# EEG Electrodes and Where to Find Them: Automated Localization From 3D Scans

**DOI:** 10.1101/2024.06.27.600334

**Authors:** Mats Tveter, Thomas Tveitstøl, Tønnes Nygaard, Ana S. Pérez T, Shrikanth Kulashekhar, Ricardo Bruña, Hugo L. Hammer, Christoffer Hatlestad-Hall, Ira R. J. Hebold Haraldsen

**Author notes:** Shared last authorship (Christoffer Hatlestad-Hall, Ira R. J. Hebold Haraldsen).

## Abstract

**Objective:** The accurate localization of electroencephalography (EEG) electrode positions is crucial for accurate source localization. Recent advancements have proposed alternatives to labor-intensive, manual methods for spatial localization of the electrodes, employing technologies such as 3D scanning and laser scanning. These novel approaches often integrate Magnetic Resonance Imaging (MRI) as part of the pipeline in localizing the electrodes. The limited global availability of MRI data restricts its use as a standard modality in several clinical scenarios. This limitation restricts the use of these advanced methods.

**Approach:** In this paper, we present a novel, versatile approach that utilizes 3D scans to localize EEG electrode positions with high accuracy. Importantly, while our method can be integrated with MRI data if available, it is specifically designed to be highly effective even in the absence of MRI, thus expanding the potential for advanced EEG analysis in various resource-limited settings. Our solution implements a two-tiered approach involving landmark/fiducials localization and electrode localization, creating an end-to-end framework.

**Main results:** The efficacy and robustness of our approach have been validated on an extensive dataset containing over 400 3D scans from 278 subjects. The framework identifies pre-auricular points and achieves correct electrode positioning accuracy in the range of 85.7% to 91.0%. Additionally, our framework includes a validation tool that permits manual adjustments and visual validation if required.

**Significance:** This study represents, to the best of the authors’ knowledge, the first validation of such a method on a substantial dataset, thus ensuring the robustness and generalizability of our innovative approach. Our findings focus on developing a solution that facilitates source localization, contributing to the critical discussion on balancing cost effectiveness with methodological accuracy to promote wider adoption in both research and clinical settings.

## 1. Introduction

Electroencephalography (EEG) employs electrodes of high sensitivity to record neuronal electrical activity at the scalp surface [1], providing crucial insight into the temporal dynamics of the brain’s neuronal mechanisms. Each EEG signal comprises a complex summation of electrical potentials induced by neuronal information transmission combined with various forms of noise and artifacts. Moreover, the EEG scalp recordings are also inherently transformed by the volume conduction effect [2], i.e., mixing the electrical currents that are conducted from their neuronal origins through organic tissues, including brain matter, cerebrospinal fluid, skull, and skin. As a result, the localization capacity of EEG signal patterns in the brain is limited. However, during the past two decades, considerable effort has been made to mitigate this challenge. One such innovation is the biophysical modelling of the head and its conductivity properties to estimate EEG signals at the neuronal (’source’) level given an individual’s anatomy. Such techniques are typically referred to as source reconstruction [3] or electrical source imaging, and are generally used to solve the inverse problem of inferring the neural origins of signals observed at the scalp (’sensor’) level. Collectively, these developments are crucial for improving the spatial resolution of EEG data, and consequently the potential of EEG in understanding brain function, e.g., in the context of connectivity [4] and pathology localization [5].

The accuracy with which source level signals can be estimated is affected by several factors, including inherent EEG signal properties such as electrode density [6], and source and head modeling errors [7]. Importantly, recent developments have paved the way for constructing head models based on realistic parameters, in contrast to analytical solutions such as the simplified spherical head model, leading to significantly more accurate representation of head geometry [8]. In particular, the accurate modeling of MRI-derived variations in skull thickness across different regions and between individuals, and the non-spherical shape of the head, are crucial to obtain high-quality source estimations [9]. The integration of these elements necessitates a model that accounts for the sensor positions relative to the estimated dipoles, which naturally differ among individuals. Consequently, the precision of individual electrode placements is therefore crucial for accurate source localization [10], validated by evidence showing that methods can precisely identify Brodmann areas with a variability of 1 cm at the sensor level [11]. Considering the brain’s complex anatomy and functional wiring, even slight errors in source localization may lead to significant misinterpretations of functional neuroimaging data.

Several methods for EEG electrode localization encompass manual, semi-automated, and automated approaches. The manually-operated Polhemus FASTRAK digitizer, often regarded as the gold standard, excels due to its precise 3D spatial registration capabilities. In contrast, manual localization software solutions, such as Fieldtrip [12], provide cost-effective alternatives despite being time-and labor-intensive. Recently, photogrammetric techniques have emerged as viable alternatives, striking a balance between accuracy and efficiency [13, 14, 15]. Additionally, innovative methods utilizing 3D scanners and lasers offer promising low-cost options [16, 17, 18, 19]. The Spot3D toolbox exemplifies the efficacy of 3D scanner technology by integrating 3D scans with individual MRI images to accurately localize electrode positions across diverse equipment [20].

While these innovative methods show considerable potential, their validation in large-scale datasets is notably still lacking. To rigorously evaluate performance across inter-individual differences, variability in data quality, and the overall robustness of methods, it is essential to use large and diverse datasets of 3D scans. This approach ensures that real-world evidence is incorporated into the evaluation. In the current study, we introduce an automated algorithm for end-to-end localization and identification of common head landmarks and high-density EEG electrodes using minimally processed 3D scans as input. The algorithm utilizes segmentation, mesh operations and alignment algorithms in its computational process. The output is notably flexible, designed to be potentially utilized with or without MRI. Previous work from our group has demonstrated the utility of combining 3D scan data and MRI data for head model construction [4]. Moreover, the developed method is subjected to the first extensive validation of an electrode localization algorithm conducted with a large dataset. This validation is critical for evaluating the applicability and efficiency of the proposed algorithm in real-world settings. Significantly, this work strives to balance cost-effectiveness and accuracy in localizing head landmarks and EEG electrode positions, thereby facilitating more accessible application of source reconstruction techniques for EEG data.

## 2. Methods and materials

### 2.1. 3D scan data

#### 2.1.1. Dataset

The 3D dataset was collected at Oslo University Hospital as part of the AI-Mind project data collection [21], using a standard operating procedure (SOP) to ensure uniformity across different data collection sites in the project. The data was generated from ANT Neuro 126 Electrode EEG Caps in two sizes: medium (red) and large (blue) [22]. The clinical dataset comprised 278 individuals aged 60-80, recorded under varying conditions of indoor lighting and seasons. This diversity introduces a wide range of edge cases, requiring robustness of the developed method. Following the SOP, the EEG cap was placed on the head, and three white markers were positioned at the landmarks: nasion and left and right pre-auricular points (L/RPA) for automated localization. Subjects, seated with faces obscured by latex-gloved hands and face masks to maintain some anonymity, were scanned with a Structure Sensor Mark II 3D scanner mounted on a 7th generation iPad, using the Scanner XRPro (3.0.3) software, starting from the front and moving around to capture the 3D scan. Crucially, the SOP defines the assumptions essential for the algorithm and its components to operate as described in this study. The pipeline requires a mesh with an overlaid texture, which facilitates the coloring of vertices essential for electrode localization. For automatic landmark localization, white markers and frontal initiation of scans are crucial. However, if that is not present, the method allows for manual placement. The electrode localization algorithm depends on landmarks—either automatically localized or manually positioned—and a template file of the cap for labeling. Moreover, scan quality and lighting substantially affected the solution’s performance. Prior to analysis, scans exhibiting inadequate data quality were excluded from the dataset. Exclusion criteria included missing mesh parts, overlapping ‘ghost’ electrodes, excessive noise, and generally poor scan quality, characterized by low resolution or blurry sections. Additionally, scans that did not comply with the SOP were excluded, for example, those with hands covering the frontal electrodes or exhibiting significant rotational irregularities. No post-processing was performed.

The complete dataset consist of 316 scans with a large (L) cap and 112 scans with a medium (M) cap. For the development and prototyping of the algorithm, 12 L cap scans were selected to reflect the significant variability typical in clinical datasets. In addition, 12 M cap scans were utilized for continous validation to confirm the method’s efficacy across both types of scans. This allocation left 304 L cap and 100 M cap scans solely for validation purposes, which were not used in the development process. Data acquisition was conducted over multiple sessions, leading to the repeated appearance of some individuals in the validation dataset: 176 subjects appeared once, 78 subjects twice, and 24 subjects three times. However, intra-subject variability, even when using the same cap, presents a challenge [23]. Consequently, data from each session must be considered independently to adequately account for this variability. In some instances, landmarks were either incorrectly positioned or completely missing, reducing the available scans for validating landmark localization to a total of 380 scans, from both M and L caps.

Ethical approval for the study was granted by the Regional Committees for Medical Research Ethics (204084). In this study, only data from the Norwegian cohort were considered. For the full ethical statement regarding the complete dataset, see the AI-Mind protocol paper [21]. All adult participants provided written informed consent to participate in this study.

#### 2.1.2. Electrode positions templates

The initial template for electrode positions was obtained from the equipment manufacturer, ANT Neuro. The PO9 and PO10 electrodes were manually adjusted by shifting them approximately 1 cm upwards to correct their initial misalignment with the cap, based on a comparison with standard positions and real-world electrode placement.

To improve algorithmic accuracy, the difference between manually localized electrodes and the automated solution, a revised template was developed based on the average positions of electrodes from 13 L cap and 8 M cap 3D scans, excluded from development and validation. The positions were manually localized using the manual adjustments option in the pipeline. This approach was predicated on the hypothesis that aligning the template more closely with the empirical data would enhance algorithm performance.

### 2.2. Landmark and electrode localization

The workflow of the automated algorithm is illustrated in Figure 1, with each component of the workflow identified by letters to facilitate cross-referencing in the subsequent detailed descriptions of each step.

**Figure 1.**
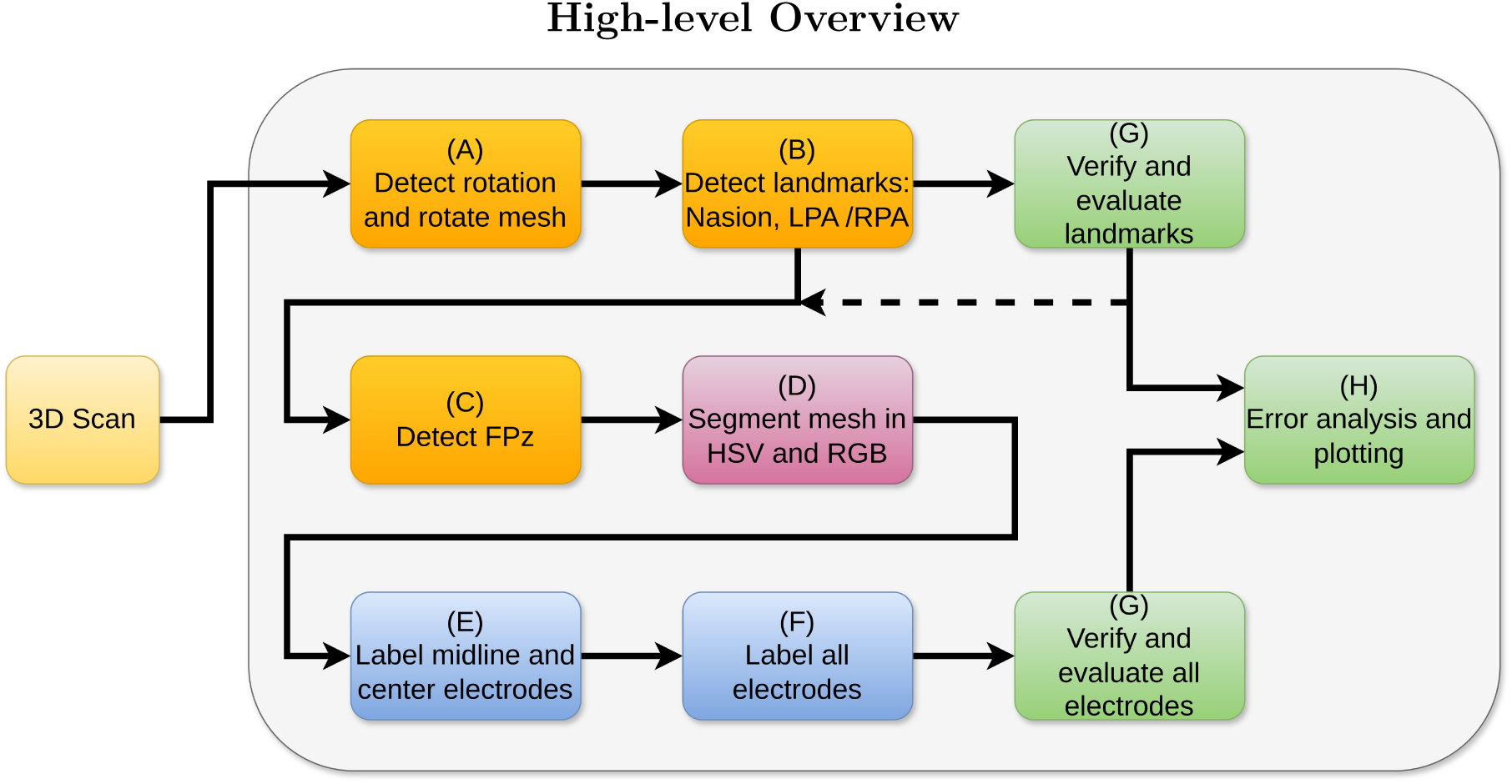
Workflow Diagram of the Automatic Algorithm. Letters within the figure are utilized as references to specific components when discussing the procedure in the text. A dashed line indicates the path followed in this study, where landmarks were validated prior to their use in electrode localization. Figure abbreviations: LPA/RPA = Right/Left Pre-Auricular; HSV = Hue, Saturation, and Value; RGB = Red, Green, and Blue.

#### 2.2.1. Mesh orientation and rotation (A)

The initial step in processing the 3D scan involves detecting the head and adjusting its orientation for uniform alignment. The mesh represents the captured headshape with the EEG cap. This process utilizes two types of cuboid bounding boxes: the mesh bounding box, defined as the minimal volume enclosing the entire mesh, and the scanning bounding box, which pertains to the spatial limits set during 3D scan acquisition.

##### Orientation detection

*1) Mesh bounding box measurement:* The procedure starts by moving 8 cm inward from the bounding box perimeter towards the center. Then, the maximal and minimal values along the remaining two axes are measured to determine the shortest distance, indicating the head’s position. *2) Planar analysis of the scanning bounding box:* The scanning bounding box often intersects the upper body, creating a plane where many vertices are closely aligned. Larger distances between these vertices typically mark the lower upper body, indicating the head’s location in the opposite direction. This spatial distribution aids in determining the head’s orientation.This method is less robust for head-only scans and serves as a secondary check to the Bounding Box Measurement Method, enhancing orientation accuracy and reliability.

##### Rotation detection

To eliminate irrelevant mesh sections, a 20 cm threshold was used based on the typical human head height of 20-25 cm [24]. The face location was estimated using curvature analysis. Significant curvature differences, especially between the front and back, were noted due to participants covering their faces with their hands. The use of light blue rubber gloves helped differentiate the brighter regions in the front from the darker colors in the back of the head. These methods assume the scan starts from the front, as specified in the SOP, and may be less effective if the face is slightly rotated.

#### 2.2.2. Landmark localization (B)

The landmark localization algorithm is vulnerable to noise, so a redundancy strategy is used. If primary measures fail, the algorithm sequentially implements alternatives, ensuring robust and reliable results.

##### Localizing nasion

First the center of the face is identified from the mesh bounding box. An offset of 2.5 cm is applied laterally from the center in both directions across the mesh. The nasion’s vertical position is anticipated to be within 1/3 to 2/3 of the head’s total height. This positioning narrows the mesh to essential features such as parts of the nose, hands, and the white marker indicating the nasion. Color segmentation isolates the three brightest clusters of vertices, which may include the nasion, Fpz electrode, face mask, or parts of the nose. These clusters consist of adjacent vertices with a similar white color, and that no points is part of other clusters. A logic-based approach using criteria like color brightness, height differences, and brightness comparisons between the top two clusters is employed. This method, adapted for edge cases in the development data set, improves the likelihood of correctly identifying the nasion, which may not always be the brightest.

##### Localizing left and right preauricular points (LPA/RPA)

A methodology similar to that used for the nasion is used to localize RPA and LPA. Mesh areas where these points are unlikely to appear, such as the face, back of the head, top of the head, and neck, are removed. Color segmentation is then employed to identify the brightest clusters of vertices. The top ten brightest clusters are identified to increase the chance of correctly localizing LPA and RPA. The center of these clusters serve as the starting points for a ray-tracing procedure, where a vector extends from each selected bright point toward the opposite side of the head. The landmark is identified at the nearest bright point along the vector’s path where it intersects with the mesh. If this method fails, a simpler logic-based approach is used as part of the redundancy strategy. This alternative strategy considers several factors: the total number of viable candidates, variations in height and distance among these candidates, color discrepancies, and ray-tracing techniques. The ray-tracing technique involves trying to match candidates in the same height. These factors are utilized to refine and select the final candidates. For detailed insights into these redundancy/logic-based methods, readers are directed to the evaluate_candidates function within the nasion and pre-auricular classes, found in the open-source code.

#### 2.2.3. Electrode localization and labeling (C-F)

Identified by its position relative to the nasion and color, the Fpz electrode serves as a strategic reference point for further electrode labeling and segmentation. Similar to the approach proposed by Taberna et al. (2019) [18], the localization primarily relies on the color differences between the electrode cap and the electrodes. The subsequent analysis primarily utilized the colored vertices, which were created by overlaying the texture onto the mesh’s structure.

##### Mesh removal

The next step involved the removal of all mesh elements located below the plane defined by the nasion, LPA, and RPA, as all electrodes should reside above this plane.

##### Segmentation

To increase the robustness of the approach under diverse lighting conditions, both red, green and blue (RGB) as well as hue, saturation and value (HSV) color schemes were used. For each scheme, the mesh underwent several segmentation iterations, employing a spectrum of threshold values ranging from stringent to more tolerant. This was designed to accommodate variations in scan quality and lighting conditions. The mesh was segmented based on white and gray colors, which represent the colors of the electrodes, and the vertices of the remaining mesh were set to black.

##### Clustering

The segmented areas were subjected to clustering using the DBSCAN algorithm [25], with an epsilon value of 0.05. The clusters were assessed based on the total number of vertices within each cluster, a parameter set within the range of 3-20 vertices to reflect the small size of an electrode. The centroid of each cluster was computed, serving as the proposed location for an electrode. The mesh with the number of proposed cluster centers closest to the total number of actual electrodes (126) was saved as the initial localized electrodes. This resulted in one set of initially localized electrode positions from RGB segmentation, and another from HSV segmentation.

##### Labeling and verifying midline electrodes

The subsequent phase involved labeling of the localized electrodes, starting with those aligned along the front-to-back midline (Fpz to Iz). The process began with aligning the template to localized positions in HSV and RGB spaces, followed by employing the Iterative Closest Point (ICP)[26] strategy to improve alignment. A verification process using electrode order and inter-electrode distances along the front-to-back midline assessed accuracy against the template. This confirms correct positioning for labeling or activates an estimation function for incorrect or missing electrodes, using template distance and direction for estimation. Subsequently, the color scheme requiring the fewest estimated electrodes was chosen for final electrode localization, giving priority to localized overestimated electrodes. The same methodology applied to the side-to-side midline (T7 to T8).

##### Labeling and verifying all electrodes

After labeling and verifying both midlines, these positions facilitated a final template alignment using ICP, labeling all localized points. Subsequently, a comprehensive verification process was initiated. This entailed sorting the electrodes by the count of their verified, labeled, and finalized neighboring electrodes. The expected position of each electrode was determined based on these neighbors, along with the specified distance and direction in the template. If a localized position aligned with its expected location, it was confirmed as a verified, finalized solution. Conversely, if misalignment occurred, estimation was undertaken based on neighboring electrodes, template-specified distance and direction, and finally, by identifying white vertices near the proposed solution. This procedure was executed for both HSV and RGB solutions.

##### Merging solutions and final verification

Finally, the localized electrodes from both color spaces were merged by averaging their positions, with additional adjustments for overlaps, where two differently labeled electrodes share the same position. The strategy for managing overlapping electrodes was as follows: if the overlap occurred solely in one color space, the electrode positions from the alternate color space were adopted. When overlaps were problematic in both color spaces, electrode positions were estimated from neighboring values, followed by a cross verification in both color spaces to guarantee consistency. The final step entailed a comprehensive verification of all electrodes, leveraging the surrounding, verified, and labeled electrodes for confirmation. This step ensured that every electrode was accounted for in the solution. Upon successful verification, the finalized solution was then displayed.

### 2.3. Validation (G)

The automatic localization framework for landmarks and electrodes includes a visual validation tool. While ideal for processing large datasets without manual intervention, this tool also allows users to manually validate and adjust EEG positions for smaller datasets or individual cases.

For the results presented in this paper, validation was structured into two distinct phases to enable separate validation of landmark and electrode localization accuracy. Firstly, the algorithm produced the suggested locations for the three primary landmarks, offering the possibility for manual corrections should these initial estimates prove inaccurate. Secondly, the algorithm provided solutions for all electrodes, allowing for similar manual adjustments if necessary. During validation, the electrodes were adjusted if the estimated position did not align with the center of the electrodes, specifically the black circle in the middle. The estimated circle had a radius of 2mm, representing the potential variation in error for the validated solutions. However, during analysis and plotting, the error distance was set to include electrodes within 5mm as correct, seen in other studies [15, 18] and reported as leading to negligible estimation errors [27]. This approach is also based on the argument that Brodmann areas can be accurately identified with variability of less than 1cm [11]. For the landmarks, solutions were classified as either correct or incorrect, based solely on whether adjustments were needed, without applying a “within 5mm correct” criterion.

### 2.4. Performance measures (H)

A comprehensive performance analysis was conducted to assess the algorithm’s efficacy across different head regions and electrodes.

#### Adjustment analysis

Analyzing the frequency and magnitude of adjustments made during the validation phase.

#### Identified vs. Approximated

The algorithm differentiates localized electrodes as “identified” when positioned correctly according to the template and “approximated” when localization or verification fails, requiring inference from neighboring electrodes, the predefined template, or both. This distinction facilitates analysis of the “identified” electrodes’ accuracy and the typical locations needing “approximation”. Notably, an electrode classified as “approximated” isn’t inherently incorrect; rather, it indicates that due to various factors, initial validation was unsuccessful, making approximation the preferred approach.

#### Neighbor distances

In specific regions of the cap, the distances between adjacent electrodes are small, potentially leading to wrong electrode localization.

#### Template difference

The template is essential in validating localized electrode positions and estimating undetected electrodes. A comparative analysis between the template positions and the average of the actual validated electrode placements across various subjects in the dataset will provide insights into the template’s representativeness and potential areas for improvement.

#### Outlier analysis

Outliers are here defined as errors above *Q*3 + 1.5 *∗* (*Q*3 *− Q*1), where Q3 is the upper/third quartile of data, and Q1 is the lower/first quartile of the data.

#### Time analysis

The primary objective is to develop a solution that is simple, accurate, and time-efficient. This requires analyzing key steps: 3D scan acquisition, computational processing time, and validation for accuracy and performance optimization.

### 2.5. Derived template analysis

The challenge posed by the inherent variability in head shapes among subjects when using a standard EEG cap template was identified during the initial validation stages. In response, the adoption of an average template, derived from actual head shapes and based on a small, distinct subset of the data, was proposed. This subset included 13 L and 8 M cap scans, deliberately excluded from the validation phase to preserve the integrity of the results. The template was generated by averaging the manually validated electrode positions across all subjects within this subset, facilitating a comparison of the effects of different templates on performance.

### 2.6. Implementation details

The pipeline is dependent on the following Python packages: 3D data manipulation utilized Trimesh [28] and PyVista [29]. Data handling employed NumPy [30]. Clustering via DBSCAN [25] and alignment through ICP [26] were executed using scikit-learn [31].

## 3. Results

### 3.1. Landmark identification

The automatic landmark localization algorithm accurately identified the nasion point in 55.8% of scans, the RPA in 80.0% of scans, and the LPA in 84.2% of scans (Figure 2). For the nasion, the median error—considering only non-correct, non-zero displacement values—was 36.4 mm, with outliers observed at approximately 100 mm and an extreme outlier near 200 mm. Similarly, the median errors for RPA and LPA, under the same conditions, were 37.2 mm and 39.3 mm, respectively, with outliers in both categories around 80 mm.

**Figure 2.**
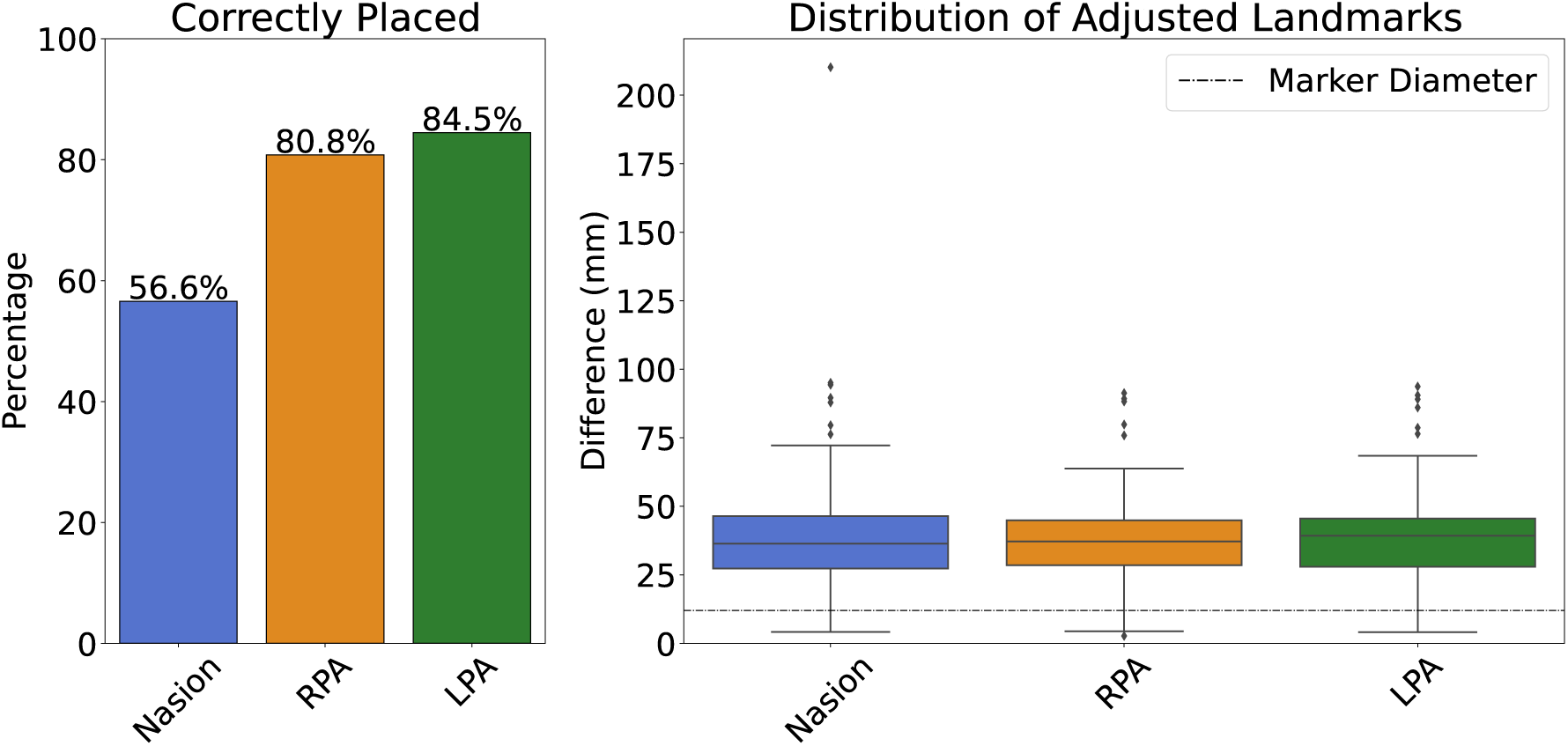
The figure illustrates the algorithm’s performance in localizing the landmark positions automatically. The histogram (left) presents the percentage of correctly localized nasion, RPA, and LPA points, with a error of less than 2 mm. The boxplot (right) displays the distribution of incorrectly placed points and their corresponding error distance in mm. The boxplot also displays the diameter of the white markers as a dotted line for reference.

### 3.2. Electrode localization

Figure 3 shows the algorithm’s performance in localizing EEG electrodes: 87.5% accuracy for L caps and 81.2% for M caps. The median error was 13.8 mm for L caps and 15.8 mm for M caps, with outliers in both cases exceeding 40mm.

**Figure 3.**
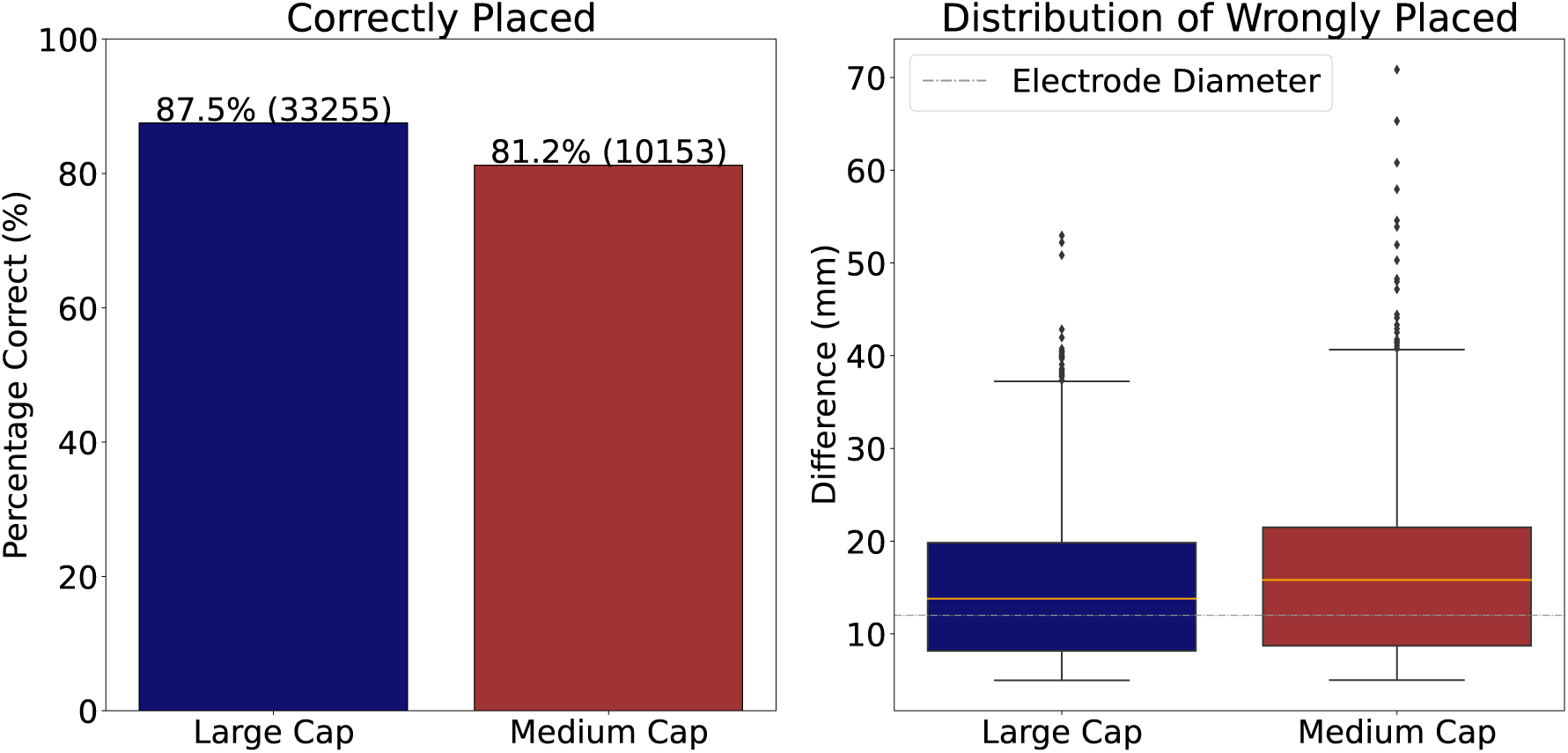
The algorithm’s efficacy in automatic EEG electrode identification for the large and medium caps. The histogram (left) displays the proportion of electrodes accurately identified by the algorithm, with an error of less than 5 mm. The boxplot (right) shows the distribution of distance errors of the wrongly placed electrodes, error above 5 mm. Additionally, the boxplot includes a gray dotted line indicating the electrode diameter for reference.

### 3.3. Performance analysis

In this subsection, the performance analysis of the method is presented. For clarity, only a subset of the full 126-channel layout is displayed in various figures. However, the complete layout is provided in the appendix, Figure Appendix A.1. When referring to channels not included in the figures in this subsection, please consult the appendix for the full layout.

#### 3.3.1. Adjustment analysis

Figure 4 highlights that outer parietal and parietal-occipital electrodes required the most frequent and significant adjustments. Key areas— AF3-AFF5h-F5, AF4-AFF6h-F6, PO4-PPO6h-P6, and PO3-PPO5h-P5—in both L and M cap sizes necessitated regular adjustments, with the M cap showing a higher need for peripheral corrections. For the L cap, adjustments had a mean frequency of 12.5% and a median of 3.29%; the M cap adjustments had a mean of 18.8% and a median of 7%. Overall, the L cap had 9 electrodes that never required adjustments, while the M cap had 22, with most of these located in the central and frontal areas. Electrodes POO9h and PPO9h were especially challenging, requiring adjustments in more than 90% of cases for both caps. In the analysis of displacement magnitude, the L cap had a mean of 1.84 mm and a median of 0.36 mm, with POO9h (15.97 mm), P9 (15.94 mm), and P10 (15.77 mm) experiencing the highest average displacements. For the M cap, the mean was 3.04 mm, the median was 0.84 mm, and the electrodes with the greatest average displacements were P9 (20.84 mm), PPO9h (19.44 mm), and P10 (19.02 mm).

**Figure 4.**
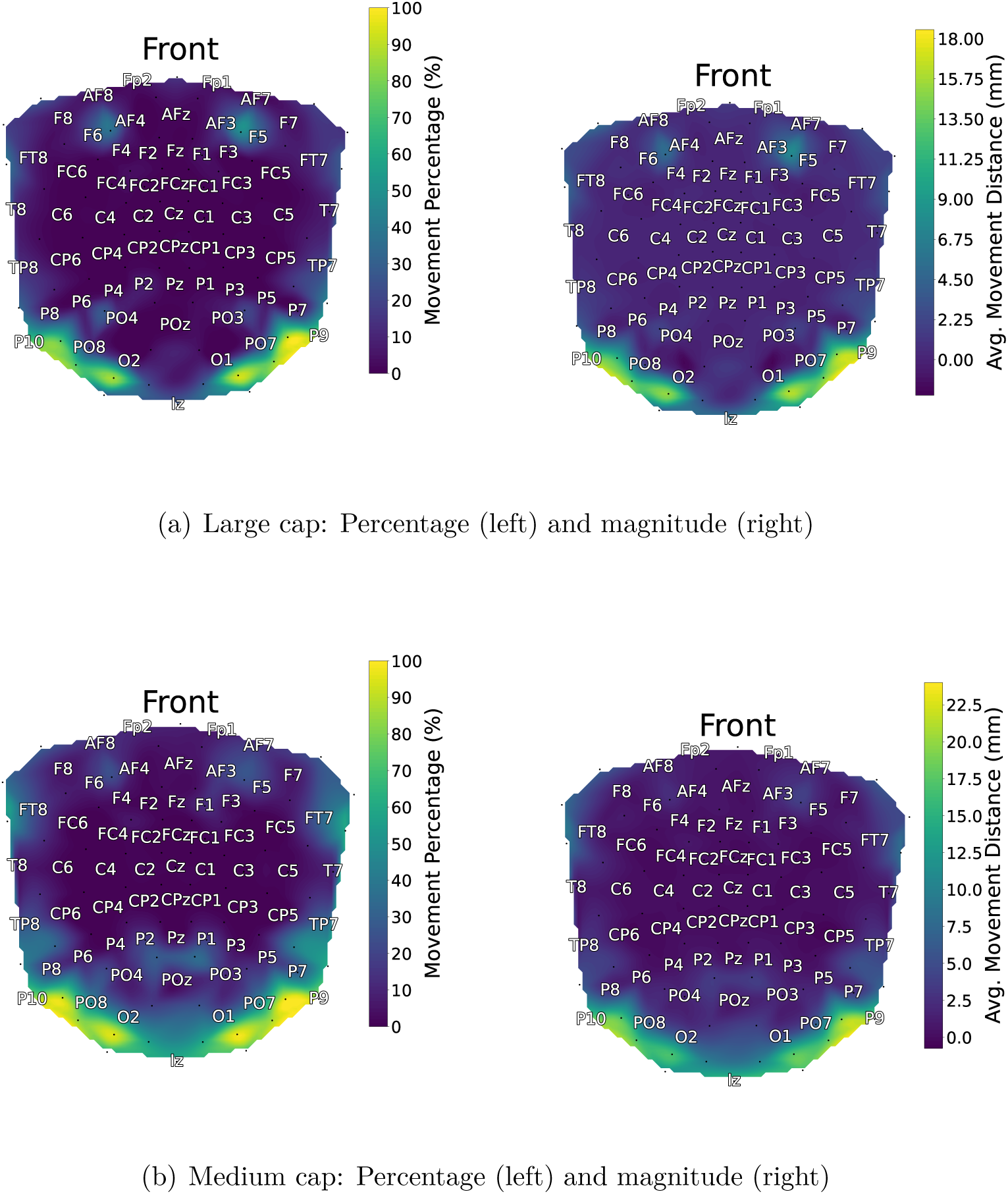
Comparison of large and medium caps: Figure 4(a) Large caps: (Left) the frequency percentage of electrode adjustments and (Right) the mean magnitude of displacement in millimeters. Figure 4(b) illustrates the equivalent metrics for medium caps. The brighter colors indicate electrodes that required more frequent adjustments or were associated with greater error distance. For the sake of visibility, only 62 of the total 126 electrodes are displayed in the figures.

#### 3.3.2. Identified vs. Approximated

Figure 5 shows that certain peripheral electrodes and specific locations (AF3, AF4, PPO6h, PPO5h, PO4, PO3) were frequently approximated in both cap sizes, with the L cap showing a higher number of identified electrodes. In the L, the most frequently approximated electrodes were P9 (97.0% of the cases), P10 (93.1%), and AF3 (92.1%), while for the M cap, F10 (99.0%), P10 (98.0%), and F9 (97.0%) were most approximated. Success rates for identified versus approximated electrodes were as follows: L caps achieved a 96.3% success rate for identified electrodes and 68.8% for approximated ones, whereas M caps recorded 93.4% for identified and 63.4% for approximated electrodes.

**Figure 5.**
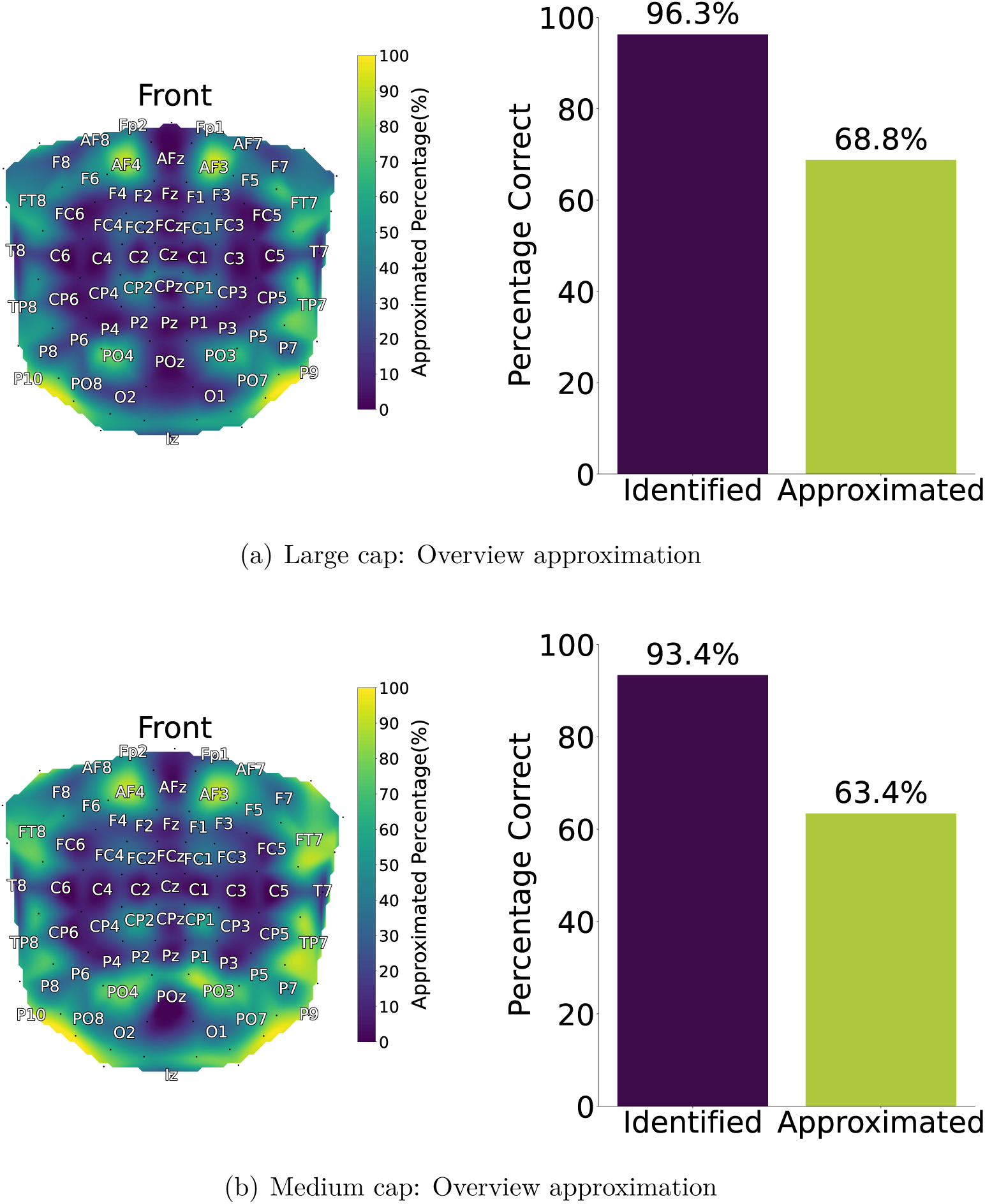
Comparison of large and medium caps: Figure 5(a) (Left) heatmap indicating the frequency of electrode approximations, with brighter colors denoting more frequent approximations, and (Right) a barplot comparing the success rate of correctly placed electrodes between identified and approximated electrodes, where the error is below 5 mm. Figure 5(b) illustrates the corresponding analyses for the medium cap. For the sake of visibility, only 62 of the total 126 electrodes are displayed in the figures.

#### 3.3.3. Closest neighbors

Figure 6 shows the distances between electrodes for both L and M caps, based on the average actual positions. Both caps exhibit similar patterns, with parietal and parietal-occipital regions, and electrodes AFF5h, AFF6h, PPO5h, and PPO6h, being consistently highlighted in the heatmaps. For both caps the closest electrodes were PPO9h, PPO10h, POO9h and POO10h.

**Figure 6.**
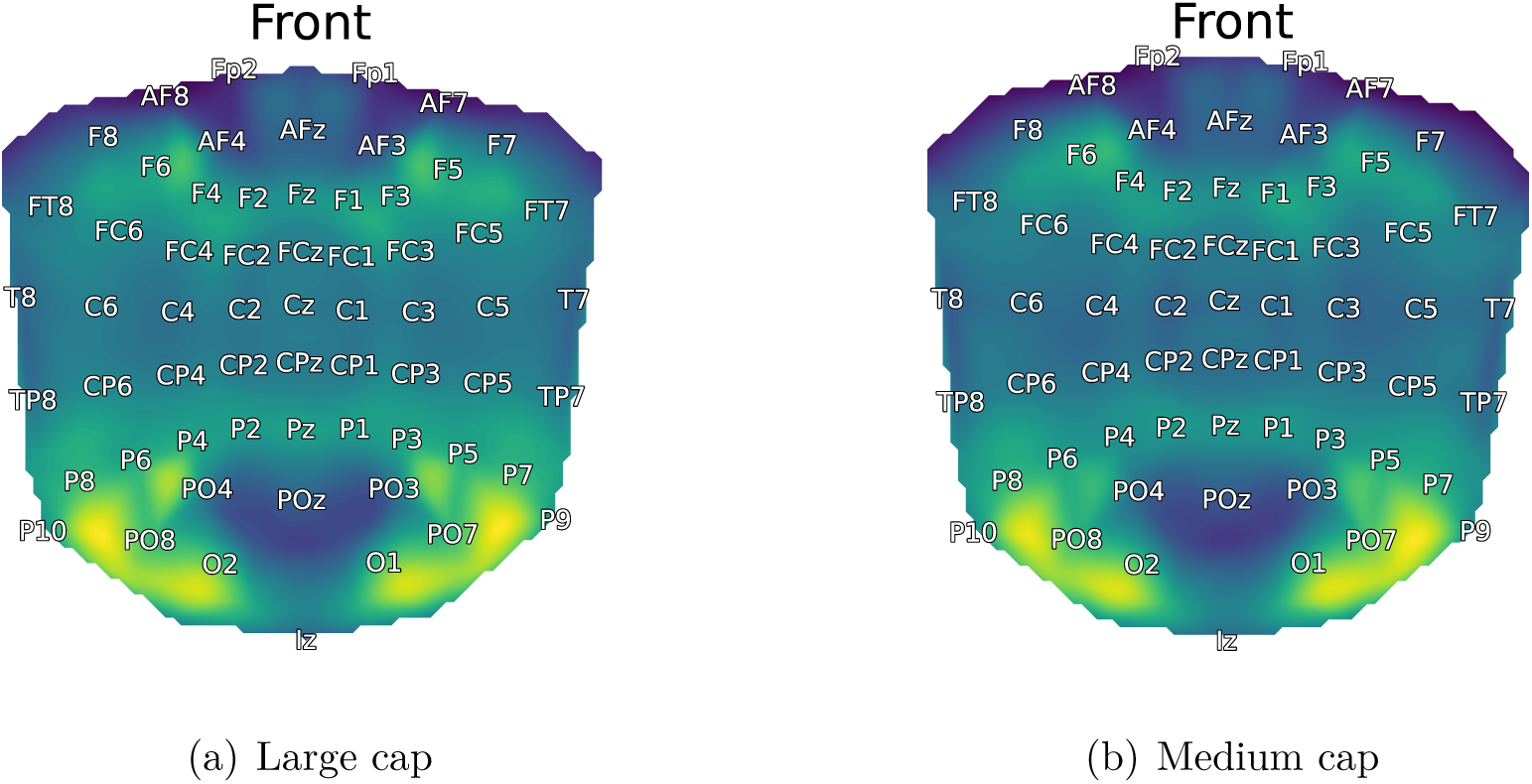
Figure 6(a) and Figure 6(b) illustrate the proximity of the three nearest neighbors, calculated from the average validated solutions for the two caps. The heatmaps display a closeness measure, the inverse of the average distances to the four closest neighbors. The brighter colors indicates closer distance to neighbors, while darker indicates more space between electrodes. For the sake of visibility, only 62 of the total 126 electrodes are displayed in the figures.

#### 3.3.4. Template vs. Average

Figure 7 demonstrates the disparities between the provided template and the averaged actual validated positions for both the L and M caps, as depicted in Figures 7(a) and 7(b), respectively. Particularly noteworthy are the most prominent differences observed in the parietal and parietal-occipital regions for both cap sizes. The averaged validated positions of the M cap exhibits nearly double the discrepancy observed in the L cap, covering a more extensive area extending from the P9-P10 line to all electrodes below this line. The mean differences were 10.64 mm for the L cap and 23.99 mm for the M cap. In the L cap, electrodes with the largest differences were P9 (25.78 mm), P009h (21.7 mm), and F10 (21.47 mm), while in the M cap, these were Iz (49.96 mm), P010 (49.29 mm), and P10 (47.58 mm).

**Figure 7.**
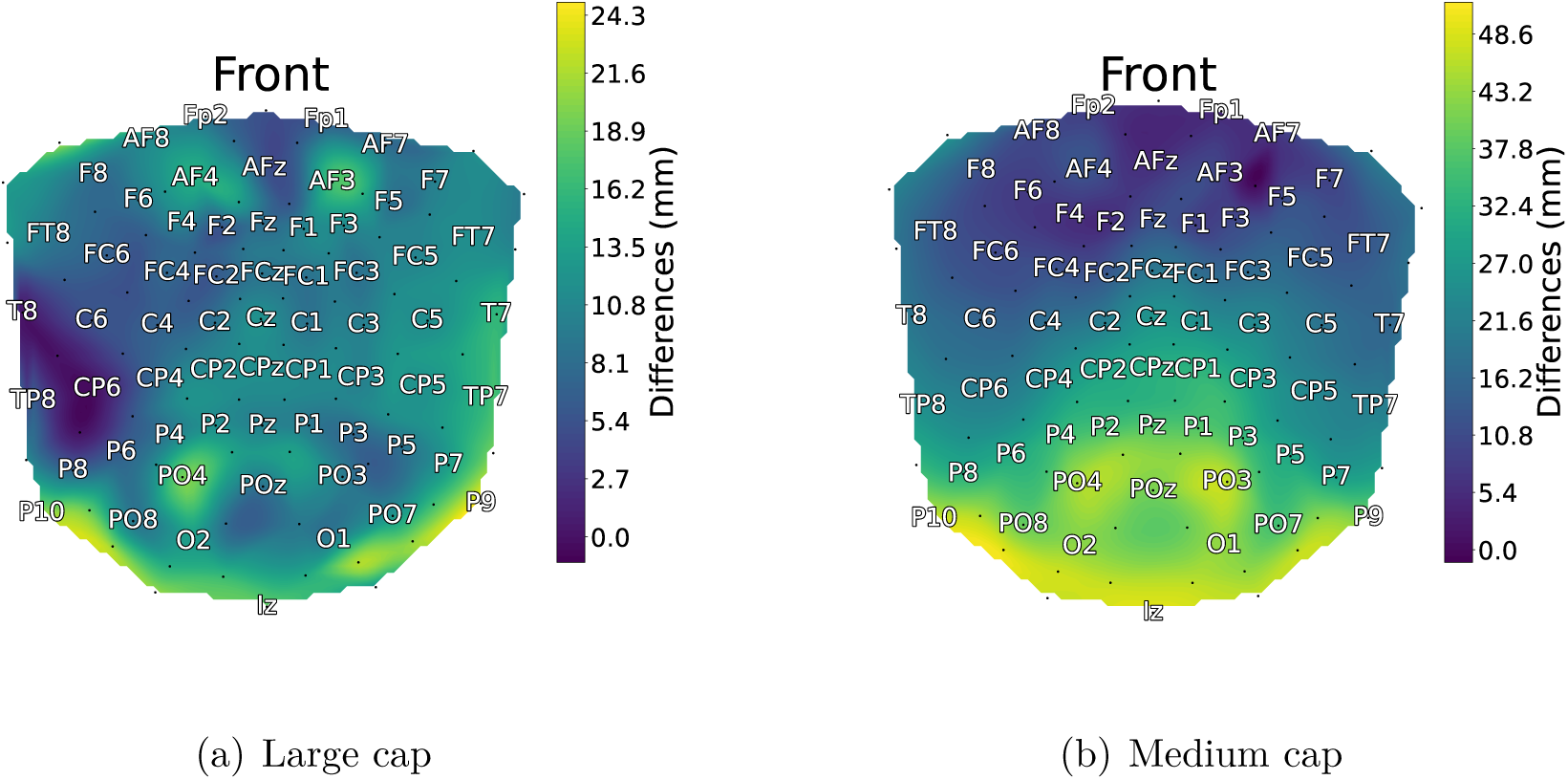
The heatmaps illustrates the differences between the average of the validated solutions for both caps and the template provided by the equipment supplier, ANT. For the sake of visibility, only 62 of the total 126 electrodes are displayed in the figures.

#### 3.3.5. Outliers

Figure 8 shows the distribution of the outliers (explained in section 2.4) for the L cap 8(a) and M cap 8(b). Notably, the identified outliers are predominantly located at the periphery of each cap.

**Figure 8.**
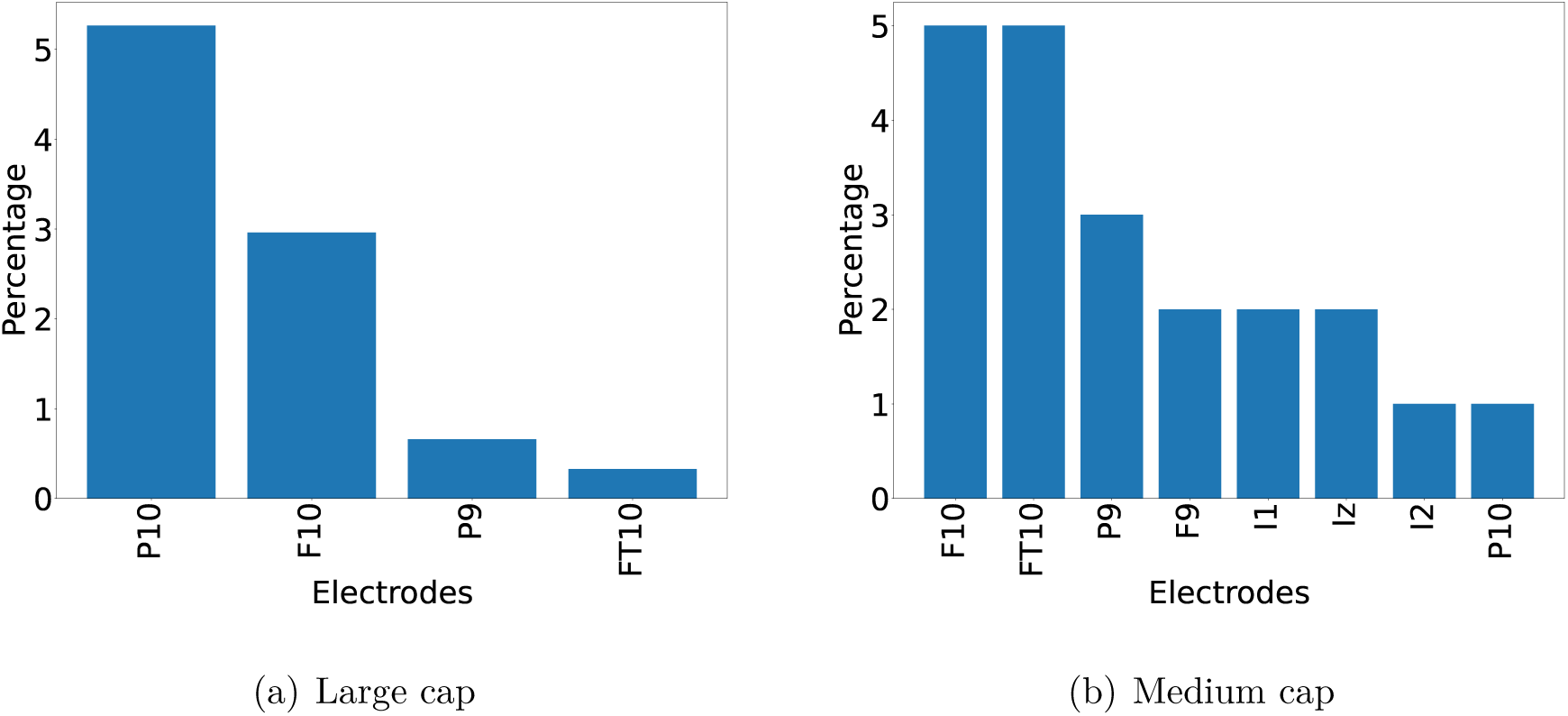
Figure presents the outliers for the different cap sizes: 8(a) illustrates the outcomes for the large cap, and 8(b) for the medium cap. The detection process identifies data points that deviate significantly from the overall dataset patterns. The x-axis represents electrodes, and the y-axis indicates the percentage of these electrodes identified as outliers. The actual positions of the electrodes can be seen in one of the previous figures, for example Figure 6.

#### 3.3.6. Timing analysis

The algorithm had a mean time of 56.2 seconds for the L caps and 57.9 seconds for M caps. The finished complete manual validation or re-adjustment had a mean time of 4.6 minutes for L caps and 5.4 minutes for M caps. This time is calculated for the original template, not the new, improved average template.

### 3.4. Template Analysis

Figure 9 demonstrates the enhanced performance achieved with the custom template, derived from actual head shapes, compared to the standard manufacturer-supplied template. Specifically, a performance improvement of 3.5 percentage points was observed for the L cap, and a 4.5 percentage points increase for the M cap. Figure 10 illustrates the improvements observed at the subject level, showing a 93.1% improvement for L caps with an average increase of 10.2 correctly localized electrodes, and a 91.0% improvement for M caps with an average increase of 13.9 correctly localized electrodes. No improvement was observed in 0.3% of the L caps, while performance worsened for 6.6% of L caps and 9.0% of M caps.

**Figure 9.**
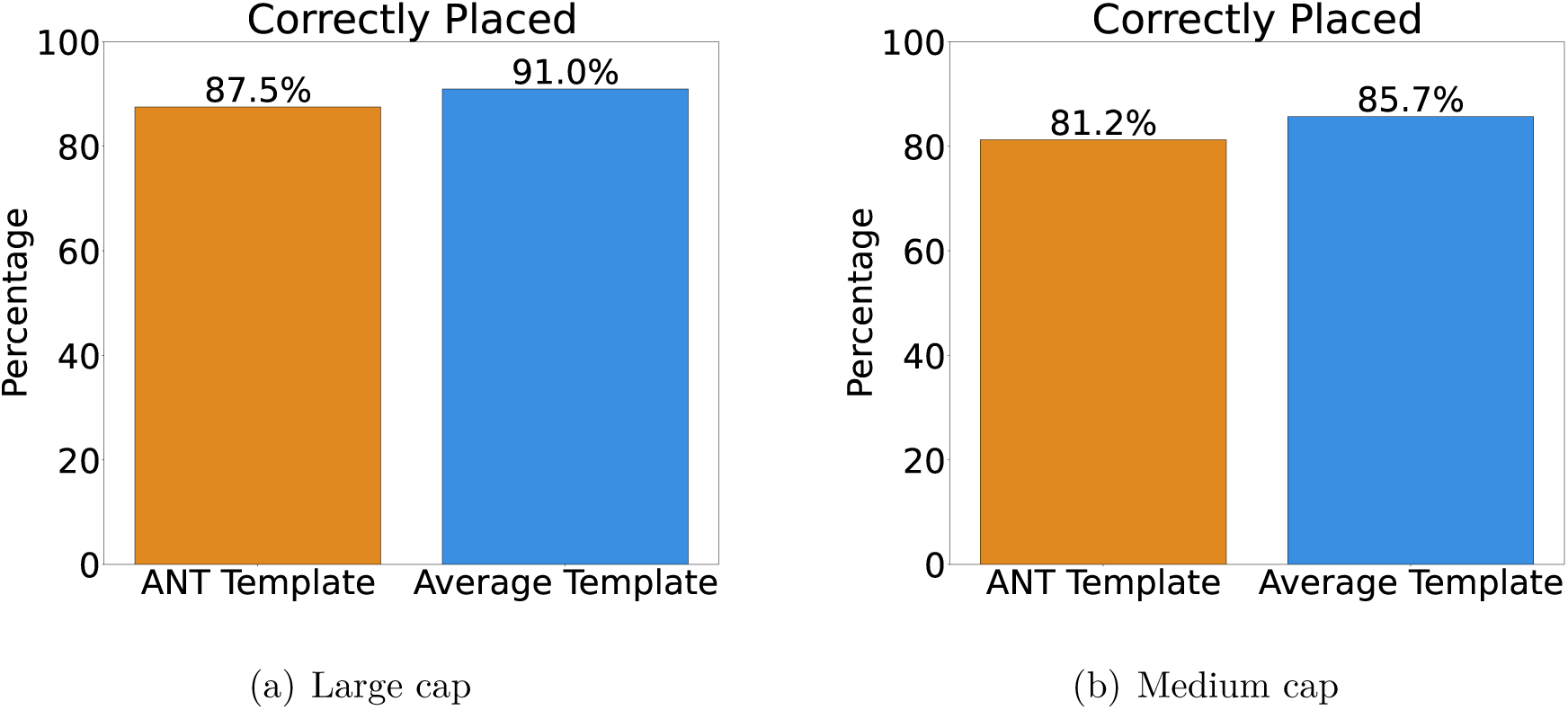
The figure compares the performance of the ANT-provided template with that of the newly developed template, distinguishing between large and medium scans in subfigures 9(a) and 9(b), respectively. Notably, the error distance is constrained to the electrode’s radius, ensuring the error measurement is within the electrode’s spatial boundaries.

**Figure 10.**
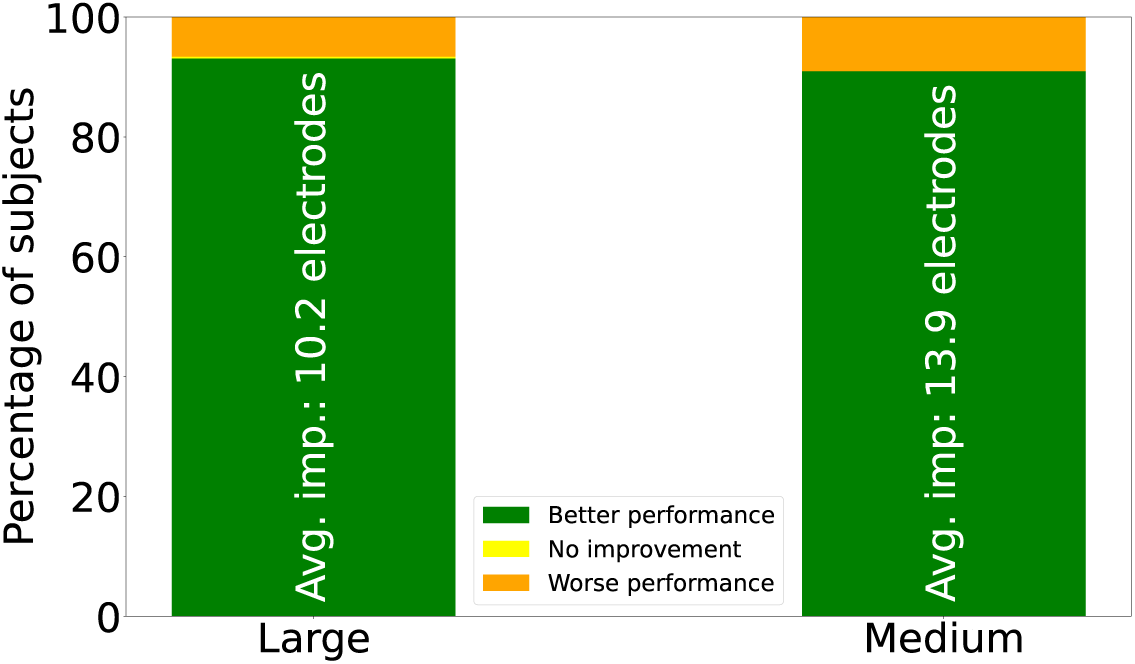
Comparative performance of the newly developed and manufacturer’s templates. The average improvement, noted within the green bars, represents the additional number of electrodes that were correctly identified when improvements occurred.

## 4. Discussion

In the current study, we propose a novel automatic method for EEG electrode localization using 3D scans, aiming to provide electrode positions for electrical source reconstruction in a cost-efficient manner. The aim was to introduce an alternative solution to currently available time-consuming or less cost-efficient methods. The resulting solution is an end-to-end automated pipeline capable of localizing a set of head landmarks and electrode positions in a high-density 126 electrode EEG cap, and validated using a large dataset.

The localization accuracies for the nasion, RPA and LPA were 55.8%, 80%, and 84.2%, respectively, with an error margin of 2mm. In the context of automatic electrode localization, absolute precision is not critical, provided that the positional suggestions do not deviate significantly enough to induce electrode shifts. However, the need for precision escalates when considering the adaptation of these solutions for future potential MRI co-registration.

The automatic electrode localization algorithm demonstrated high performance, achieving overall accuracy of 87.5% for 304 L caps and 81.2% for 100 M caps, spanning all electrodes, within a 5mm error margin.

Analyzing outliers highlights the methods’ sensitivity to the Standard Operating Procedures (SOP). Strict adherence to SOPs is crucial for ensuring data integrity and reliability, particularly in automated systems that lack human oversight. Without sufficient compliance with SOPs, there is no assurance that the data accurately represents the intended subject, such as a human head, even though algorithms may accommodate variations in rotation or shape. Discrepancies in rotation due to non-adherence to SOPs can significantly degrade the algorithm’s performance, as evidenced by outliers in nasion and RPA/LPA measurements, leading to errors in landmark identification. For example, the algorithm might mistakenly select other bright areas such as different electrodes for the nasion, and misidentify hair, electrodes, or brightly lit ear regions as landmarks for RPA/LPA. Such misidentification of landmarks can cause shifts in electrode placement during the localization process in a fully automated pipeline. Outlier analysis for electrode localization revealed that most outliers were found at the cap edges, often linked to inaccurately placed or absent landmark markings, as verified through manual checks. This frequently led to the exclusion of mesh sections containing electrodes, particularly when landmarks were positioned too high, thus removing these electrodes from the analysis.

Incorrect localization of landmarks were predominantly attributed to lighting conditions and scan quality, resulting in misidentifications in excessively illuminated areas, such as portions of the ear. The mandatory use of facemasks during data collection (due to Covid) occasionally caused the algorithm to mistake the white brim of facemasks for the nasion points. Despite this challenge, the localization of the LPA and RPA achieved good results, underscoring the robustness of our approach. For the wrongly placed electrodes a more detailed performance analysis was conducted. The algorithm was particularly good at localizing electrodes in the frontal and central regions. We achieved lower accuracy in parietal, parietal-occipital regions, and specific zones (AF3-AFF5h-F5, AF4-AFF6h-F6, PO4-PPO6h-P6, and PO3-PPO5h-P5), where electrodes frequently required adjustments. Large differences were observed in parietal and occipital areas, likely due to substantial variations between actual head shapes and the provided template, especially pronounced in M caps. The four electrode zones were identified as areas with minimal inter-electrode distances, elevating the likelihood of localization errors. Electrodes in the final solution were classified as “identified or “approximated” to provide insights, with “identified” electrodes correctly localized in 93.4% of M caps and 96.3% for L caps, contrasting with a 63.4% and 68.8% accuracy rate for “approximated” electrodes for M and L, respectively. This distinction also offers a valuable metric for assessing uncertainty and identifying regions that might require manual validation in practice.

Importantly, we observed that the suboptimal localization of electrodes was mitigated when the standard position template was substituted by a template derived from a subset of data not used in training nor validation. The accuracy increased by 3.5 percentage points in performance for L caps to 91.0% and 4.5 percentage points for M caps to 85.7% in terms of correctly placed electrodes. Investigations at the subject level revealed that over 90% of the subjects experienced an increase in performance with the new template, correctly localizing more than 10 additional electrodes on average for both cap types. However, a subset of subjects also experienced a decrease in performance, potentially due to electrode shifts, which resulted in the displacement of a large number of electrodes. Similar issues were occasionally observed with the original template, which may have been caused by various factors, including improper template fit or incorrect detection of key areas such as the midlines. Template discrepancy was also reported by Homölle et al.[19] as a potential problem. It remains possible that the accuracy increase might have been greater had the template been more finely tuned to the data, e.g., by basing it on more 3D scans. Verifying the representativeness of the external validation set concerning head shape is not conducted, as such analysis falls outside the article’s scope. Nonetheless, it is recognized that careful analysis and generation of templates could significantly enhance representativeness and applicability.

Comparing our results with those from studies such as Taberna et al. [18] and Hômolle et al. [19] is challenging, primarily due to differences in validation techniques and equipment—a complication also noted by Taberna et al. Our validation, conducted exclusively within our framework without external tools like FieldTrip or Polhemus, has occasionally resulted in accurately positioned electrodes being assigned zero values—a consequence that the mean and average measurements will misrepresent the algorithm’s true accuracy. Additionally, the inherent limitations of 3D scans, susceptible to errors, alongside variations in preprocessing between studies, complicate the evaluation further. Distinguishing whether inaccuracies arise from the 3D scans or the algorithm itself is imperative for a comprehensive assessment of our method’s efficacy, underscoring the difficulties in achieving reliable external validation. Given these complexities, external validation is outside the scope of the paper.

The data collection protocol included the use of white markers to mark the landmarks, a step that may appear counterintuitive compared to simply manually selecting these points using software. However, we justify this approach for two primary reasons. Firstly, when validating a large dataset, our method’s automatic localization proves efficient, saving time and effort in subsequent analyses. Secondly, the manual placement during cap fitting is a straightforward and quick process, integrated seamlessly into the workflow. An important aim of the currently proposed method is to minimize the time required for extracting EEG electrode positions. The acquisition time for the 3D scans was in line with previous studies, around 2 minutes [18, 19]. While the algorithm’s computational speed has yet to be optimized, e.g., by implementing parallelization techniques, the current processing time for both L and M caps is approximately 1 minute, which we consider reasonable in the context of a clinical study. The validation step showed that L caps took approximately 4.6 minutes and M caps around 5.4 minutes for both landmark and electrode validation. The potential for time optimization with the new template could arise from the reduced need for electrode adjustments.

To the best of our knowledge, this study represents the first application of a large-scale dataset with over 400 3D scans specifically for EEG electrode localization, thereby constituting a significant advancement in the field. The importance of using a comprehensive dataset to analyze results and refine methodologies cannot be overstated. Such a dataset captures the full spectrum of potential inconsistencies that might arise during data collection processes, such as the occasional misplacement of white landmark markers due to the tight schedule encountered in clinical settings. These situations can introduce outliers and anomalies in electrode localization, emphasizing the need for robust localization methods. The observed variability across different scenarios— stemming from changing data quality, lighting, subject variability, and other factors— highlights the limitations of small, highly controlled datasets. There is a recurring trend in recent studies to propose time-efficient methodologies yet validate them using small (<30), non-representative, and highly controlled datasets. While these smaller datasets may offer a small measure of precision, they often fail to represent the complexities and challenges of real-world data collection, potentially leading to an overestimation of a method’s true effectiveness. Thus, the integration of a diverse and sizable dataset is vital for a comprehensive assessment and enhancement of localization techniques. This strategic approach ensures the evaluated method’s performance is a true reflection of its capabilities in varied and realistic settings. Our robust and automatic framework, demonstrating overall good performance, is built in a modular fashion, aimed for future refinement, and offers the possibility to add functionality and tailor it to future needs. This adaptability underscores the framework’s versatility and its potential to evolve alongside emerging needs, positioning it as a dynamic and enduring solution.

### Adaptability, Limitations and Future Work

As EEG systems differ substantially in terms of electrode mounting (e.g., elastic caps, nets, fastening directly to the scalp with paste), creating a 3D scan-based electrode position identification system that captures all systems is challenging. While the development of the current method was tailored to a particular EEG system, the method’s adaptability to new datasets is partially addressed by the modular design of the framework. This modular approach facilitates the future integration of modules for other systems, file types, and scanners. Although the robustness of our automated procedure has not been evaluated beyond the specific dataset and SOP, it is important to recognize that certain components of the framework may still operate effectively even if the standard procedures are not strictly followed. In such instances, manual input of landmarks might be required, but automatic electrode localization should remain achievable. However, it is important to note that compatibility issues may arise with equipment from different suppliers or different cap types, as the method depends on detecting color differences, particularly lighter colors. Nonetheless, if there is a similar contrast between light-colored electrodes and dark cap colors, adaptation of the method could still be feasible.

This study presents certain constraints that also outline avenues for future research. The algorithm was validated with data exclusively from one laboratory, incorporating an array of photographers, subjects, seasonal variations, and lighting conditions. This evaluation was conducted solely on ANT caps, indicating potential adaptability requirements for other cap designs. Additionally, preprocessing steps such as light correction and mesh improvement were not applied, with the rationale being to challenge the algorithm with a broad spectrum of scan qualities. Introducing preprocessing steps, such as light correction and general mesh improvement strategies, could potentially reduce the complexity of the input data, which might not fully test the algorithm’s robustness in handling diverse conditions. A notable constraint of our approach is its inapplicability to retrospective data, making alternative strategies like those suggested by Martinez et al. [32], which employ automatic localization of landmarks through algorithms for eyes, ears, and facial recognition, more suitable. However, our system incorporates a validation tool that facilitates rapid corrections of incorrectly positioned landmarks and allows for manual placement as well. This feature significantly enhances the practicality and usability of our approach in clinical and research settings.

For future work, several avenues are interesting. Firstly, investigate the influence of preprocessing on performance, with an expectation of performance enhancement. Secondly, how electrode positions, determined by this method, affect the precision of source reconstruction. Furthermore, it may be relevant to conduct a cost-benefit analysis to evaluate whether the outcomes of this approach justify its adoption over more accurate but potentially costlier alternatives. Lastly, prioritizing the exploration of robust methods for comparing results between studies is essential.

## 5. Conclusion

We have developed and validated a robust framework for localizing EEG electrodes using 3D scan data, validated its effectiveness across a substantial dataset. Our framework demonstrated high efficiency, achieving a mean accuracy of over 90% in correctly localizing electrodes. The high accuracy, combined with the option for manual validation and adjustment of incorrectly detected electrodes, makes the solution highly effective. This work provides a promising avenue for future research, with the aim of enhancing the practicality and accessibility of EEG analysis in healthcare settings.

## Project Information and Contributions

### Funding

This project has received funding from the European Union’s Horizon 2020 research and innovation programme under grant agreement No 964220. This publication reflects views of the authors and the European Commission is not responsible for any use that may be made of the information it contains.

### Author Contribution

**Mats Tveter:** Software, Investigation, Methodology, Validation, Formal Analysis, Writing – Original Draft **Thomas Tveitstøl:** Investigation, Writing – Review & Editing **Tønnes Nygaard:** Supervision, Methodology, Writing – Review & Editing **Ana S. Pérez T.**: Writing – Review & Editing, **Shrikanth Kulashekhar:** Writing – Review & Editing **Ricardo Bruña:** Writing – Review & Editing **Hugo L. Hammer:** Supervision, Writing – Review & Editing **Christoffer Hatlestad-Hall:** Supervison, Conceptualization, Writing – Review & Editing **Ira R. J. Hebold Haraldsen:** Project administration, Writing – Review & Editing, Funding acquisition, Resources

### Data Availability

The data is currently unavailable due to privacy concerns related to the 3D scans. However, a fully anonymized version of the data will be made available in the future, alongside the rest of the AI-Mind dataset. The code in its entirety is available at Github^‡^.

## Appendix A. Electrode layout

**Figure Appendix A.1.**
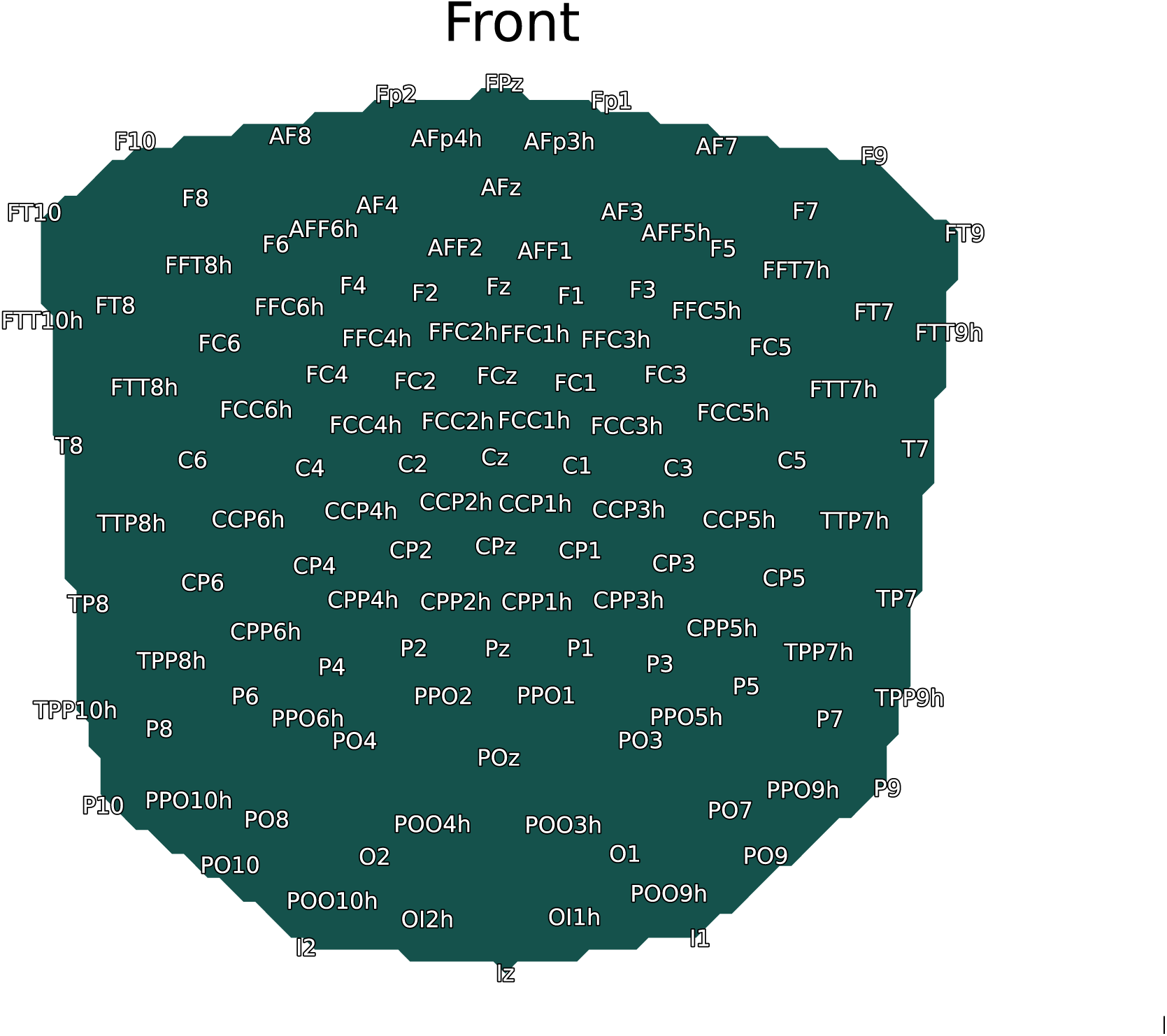
Figure displaying the layout of the 126 channel ANT electrode cap.

## Footnotes

‡ https://github.com/matstveter/EegElectrodeLocalizer

